# miR6236, a microRNA suppressed by the anisotropic surface topography, regulates neuronal development and regeneration

**DOI:** 10.1101/2020.07.12.196675

**Authors:** Yi-Ju Chen, Yung-An Huang, Chris T. Ho, Jinn-Moon Yang, Jui-I Chao, Ming-Chia Li, Eric Hwang

**Affiliations:** Institute of Molecular Medicine and Bioengineering, National Chiao Tung University, Hsinchu, Taiwan; Department of Biological Science and Technology, National Chiao Tung University, Hsinchu, Taiwan; Institute of Bioinformatics and Systems Biology, National Chiao Tung University, Hsinchu, Taiwan; Center for Intelligent Drug Systems and Smart Bio-devices (IDS^2^B), National Chiao Tung University, Hsinchu, Taiwan

## Abstract

It has been well studied that the surface topography affects the growth and development of neurons. However, the precise mechanism that the surface topography leads to cellular changes remains unknown. In this study, we created an anisotropic surface using nanodiamonds and discovered this surface topography accelerates the development of primary neurons from both the central and peripheral nervous systems. Using RNA sequencing technology, a previously uncharacterized microRNA (miR6236) was found to exhibit significant and the most substantial decrease when neurons are cultured on this nanodiamond surface. Gain- and loss-of-function assays confirm that miR6236 is the predominant molecule responsible for converting the surface topography into biological responses. We further demonstrate that the depletion of miR6236 can enhance neuroregeneration on inhibitory substrate, uncovering its therapeutic potential for promoting central nervous system regeneration.

## Introduction

All vertebrate animals interact with their environment using a complex network of interconnected cells called the nervous system. At the center of this interconnected network of cells is the neuron, a special type of cell with excessively long and elaborated cellular protrusions. Neurons acquire this sophisticated morphology via a serial of stereotypical steps during their development. Using hippocampal neurons cultured *in vitro*, it has been documented that the morphological development follows some stereotypical steps including the sprouting of the neurite (i.e. the premature axon or dendrite), the elongation of the neurite, the polarization of the axon, the formation of the dendrite, the elaboration of branches, as well as the formation of synapses (Dotti et al., 1988).

Both intrinsic and extrinsic factors influence the morphogenetic process of neurons (Cheng and Poo, 2012; Dong et al., 2015). It has been shown that various extrinsic factors contribute to the morphological development, including the biochemical and physical microenvironment. Biochemical microenvironment has been known to play important roles in regulating the processes of axon guidance (Dickson, 2002), axonal/dendritic development (Jan and Jan, 2003), branching (Jan and Jan, 2010; McAllister, 2000), and many others. During neuronal development, the growth cone senses biochemical factors in the environment and activates Rho GTPases to regulate the actin cytoskeleton, which results in morphological changes (Govek et al., 2005; Lowery and Van Vactor, 2009). In addition to regulating Rho GTPases, it has also been shown that biochemical microenvironment regulates the expression of microRNAs (miRNAs) (Vo et al., 2005). miRNAs are small non-coding RNAs consist of ~22 nucleotides and mediate gene silencing. They are loaded into the miRNA-induced silencing complex (miRISC), which facilitates translation repression or mRNA cleavage (Jonas and Izaurralde, 2015). The targeting of miRISC is mediated by the base pairing between the seed region of the miRNA and the 3’ UTR of its target mRNA (Krol et al., 2010).

Furthermore, accumulating evidence indicates that not only biochemical but also physical microenvironment plays important roles in the morphological development of neurons. The brain is a complex three-dimensional structure with heterogeneous stiffness and surface topography (Amunts and Zilles, 2015; Koser et al., 2016). For the effect of stiffness on neurons, Flanagan *et al.* first reported that primary neurons extended longer neurites and developed higher complexity on a soft surface compare to the rigid one (Flanagan et al., 2002). In the past 2 decades, accumulating evidences demonstrate that neurons can sense and respond to the stiffness in their surrounding environment (Franze et al., 2013). For the effect of surface topography on neurons, the field is filled with studies using numerous materials with various surface topography, these include nanoscopic roughness [e.g. nanodiamonds (NDs), beads, and others (Brunetti et al., 2010; Cyster et al., 2004; Edgington et al., 2013; Huang et al., 2014; Kang et al., 2012b; Kang et al., 2014; Khan et al., 2005; Thalhammer et al., 2010)], anisotropic pattern [e.g. ridges, grooves, gratings, and electrospun fibers (Baranes et al., 2012; Clark et al., 1991; Huang et al., 2018; Johansson et al., 2006; Kang et al., 2012a; Kang et al., 2011; Nagata et al., 1993; Nikkhah et al., 2012; Patel et al., 2007; Rajnicek and McCaig, 1997; Schnell et al., 2007; Tonazzini et al., 2014; Webb et al., 1995; Xie et al., 2009; Yang et al., 2005)], and isotropic pattern [e.g. pillars, nanotubes, and columns (Dowell-Mesfin et al., 2004; Hanson et al., 2009; Hu et al., 2004; Mattson et al., 2000; Park et al., 2016; Roberts et al., 2014; Xie et al., 2010)]. More comprehensive reviews about the surface topography on neuronal morphogenesis have been published elsewhere (Hoffman-Kim et al., 2010; Simitzi et al., 2017). These studies demonstrate that neurons can sense and respond to their physical microenvironment. However, the biological mechanism of translating topographic features into the morphological response remains largely unknown.

In this study, we aim to elucidate the cellular response elicited by the surface topography on neuronal development. A surface with irregular nanotopography was created using 3~5 nm NDs, and it promotes neuronal attachment, neurite elongation, as well as axon polarization. To understand the biological response elicited by the ND-coated surface (hereafter referred to as ND surface), RNAseq was utilized to examine the transcriptomic differences in neurons cultured on flat or ND surface. Surprisingly, a single miRNA (miR6236) essentially accounts for the transcriptomic change between the two surfaces. miR6236 is the most abundant miRNA in early stage neurons on a flat surface. Its expression is significantly reduced on ND surface. Loss- and gain-of-function analyses confirm that miR6236 is predominantly if not entirely responsible for the developmental changes on ND surface. Given that miR6236 negatively impacts neuronal development, we tested the possibility to promote neuronal regeneration by depleting miR6236. Using dissociated hippocampal neurons cultured on an injury-mimicking environment, this therapeutic potential of miR6236 was confirmed. These results provide the first connection between the surface topography and miRNA expression, and discover the therapeutic potential of this miRNA on neuronal regeneration.

## Results

### Nanodiamond surface promotes neuronal attachment and neurite elongation

We have previously shown that NDs do not induce cytotoxicity in dissociated primary neurons *in vitro*, yet adding NDs to the culture after neurons are attached to the substrate compromises neurite elongation due to the vibration of NDs in solution (Huang et al., 2014). Here, we investigated whether pre-coating the culture surface with NDs (which makes NDs immobile) can affect the extension of neurites. Our coating procedure generated a rugged surface topography as seen under scanning electron microscope (**Figure S1**). At lower density (9.4 or 37.5 μg/cm^2^), most NDs exist in small clusters, while some large clusters exceeding 100 nm in diameter can also be detected (**Figure S1A-D**). At higher density (150 μg/cm^2^), most NDs appear as an uneven covering on the surface (**Figure S1E-F**).

We first examined the effect of immobile NDs on neuronal attachment. Dissociated neurons are historically cultured on surface coated with positively changed polypeptides (e.g. poly-lysine) to facilitate their attachment (Kaech and Banker, 2006). To our surprise, ND surface enables the attachment of dissociated hippocampal neurons as good as the standard flat surface coated with poly-L-lysine (hereafter referred to as PLL surface) (**Figure 1A-B**). Between coating density 4.7 ~ 37.5 μg/cm^2^, hippocampal neurons cultured on ND surface sprout longer neurites than those cultured on PLL-coated surface (hereafter referred to as PLL surface) (**Figure 1C**). It has been shown that neurites from dissociated mouse neurons tend to form bundles when cultured on PLL surface in normoxic condition (Chen et al., 2011; Peacock et al., 1979). Interestingly, ND surface significantly reduces this bundling phenomenon compared to PLL surface (**Figure 1D**). Because the automatic neurite quantification software we used are unable to distinguish a single neurite from a group of bundled neurites, it is a possibility that the longer neurite length on ND surface was the result of less neurite bundling. To test this possibility, we seeded neurons at a low density which essentially eliminates neurite fasciculation. Manual neurite tracing was performed in addition to automatic tracing to distinguish single neurites from bundled ones (**Figure S2**). Neurons cultured on ND surface still produce longer neurites under this condition, demonstrating that ND surface indeed promotes neurite elongation.

**Figure 1.**
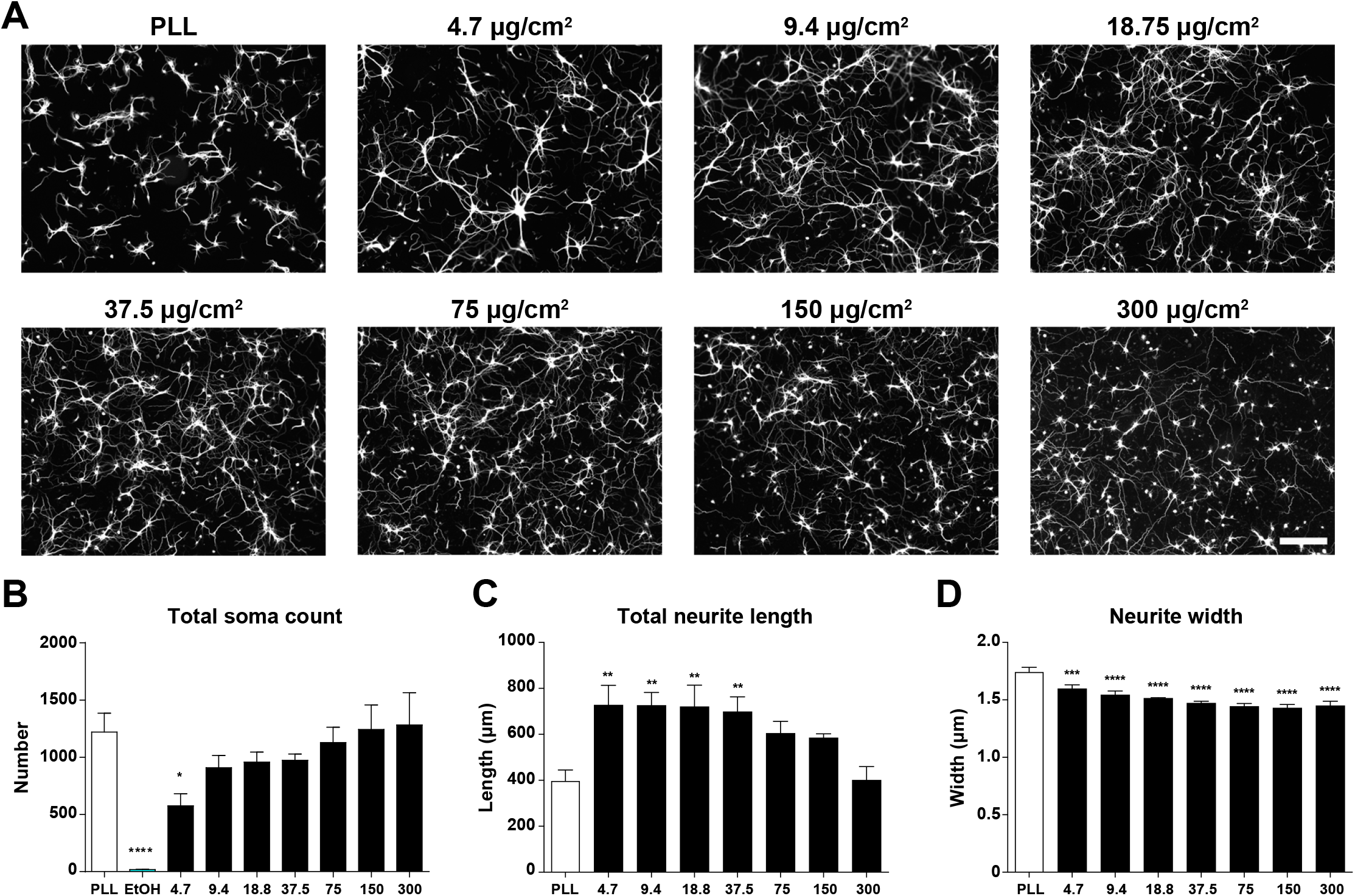
Nanodiamond surface promotes the attachment of hippocampal neurons in a dosage-dependent manner. (A) Representative images of 3DIV dissociated hippocampal neurons cultured on surfaces coated with various densities of NDs. Neurons are identified by immunofluorescence staining with the antibody against β-III-tubulin. All images have the same scale and the scale bar represents 200 μm. Quantifications of total soma count in a 13.91 mm^2^ area (B), total neurite length per neuron (C), and average neurite width (D). * *p*<0.05, ** *p*<0.01, *** *p*<0.001, **** *p*<0.0001, one-way ANOVA followed by Dunnett’s post-hoc analysis against the control group (PLL). All bar graphs are expressed as mean and SEM from 3 independent repeats.

In addition to the principal neurons from the central nervous system, sensory neurons from peripheral nervous system (i.e. dorsal root ganglion or DRG neurons) also respond to ND surface in a similar manner. ND surface facilitates the attachment of DRG neurons and enables the outgrowth of axons comparable to that on PLL surface (**Figure S3**).

### Nanodiamond surface accelerates neuronal polarization

It has been shown that the surface made from submicrometer self-assembled silica beads accelerates the development of hippocampal neurons (Kang et al., 2012b). Given the ND surface possess topography in the submicrometer scale, we were curious whether it can also accelerate neuronal development. The hippocampal neuron developmental process *in vitro* can be classified into several different stages (Dotti et al., 1988). In stage 1, neurons extend cellular protrusions that are rich in actin cytoskeleton (e.g. lamellipodia and filopodia). In stage 2, neurons start to sprout microtubule-based protrusions called neurites. In stage 3, one of the neurites exhibits accelerated outgrowth and expresses axon-specific proteins. Neurons at this stage is also known as polarized neurons. As a result, the progression of neuronal polarization can be quantified based on the proportion of neurons entering stage 3 at a given time. Dissociated hippocampal neurons were cultured on PLL or ND surface for 40 hours and immunofluorescence stained with axon- and dendrite-specific markers to determine whether it has entered stage 3 (**Figure 2A and C**). While less than 40% of the hippocampal neurons cultured on PLL surface enter stage 3 at this time, a significantly higher proportion (>75%) of neurons on ND surface are polarized (**Figure 2C**). Interestingly, a significantly higher percentage of polarized neurons possesses multiple axons (i.e. a neuron with 2 or more axons) when cultured on ND surface than on PLL surface (**Figure 2B and 2D**). These data demonstrate that ND surface not only accelerates neuron polarization, but also induces the formation of multiple axons.

**Figure 2.**
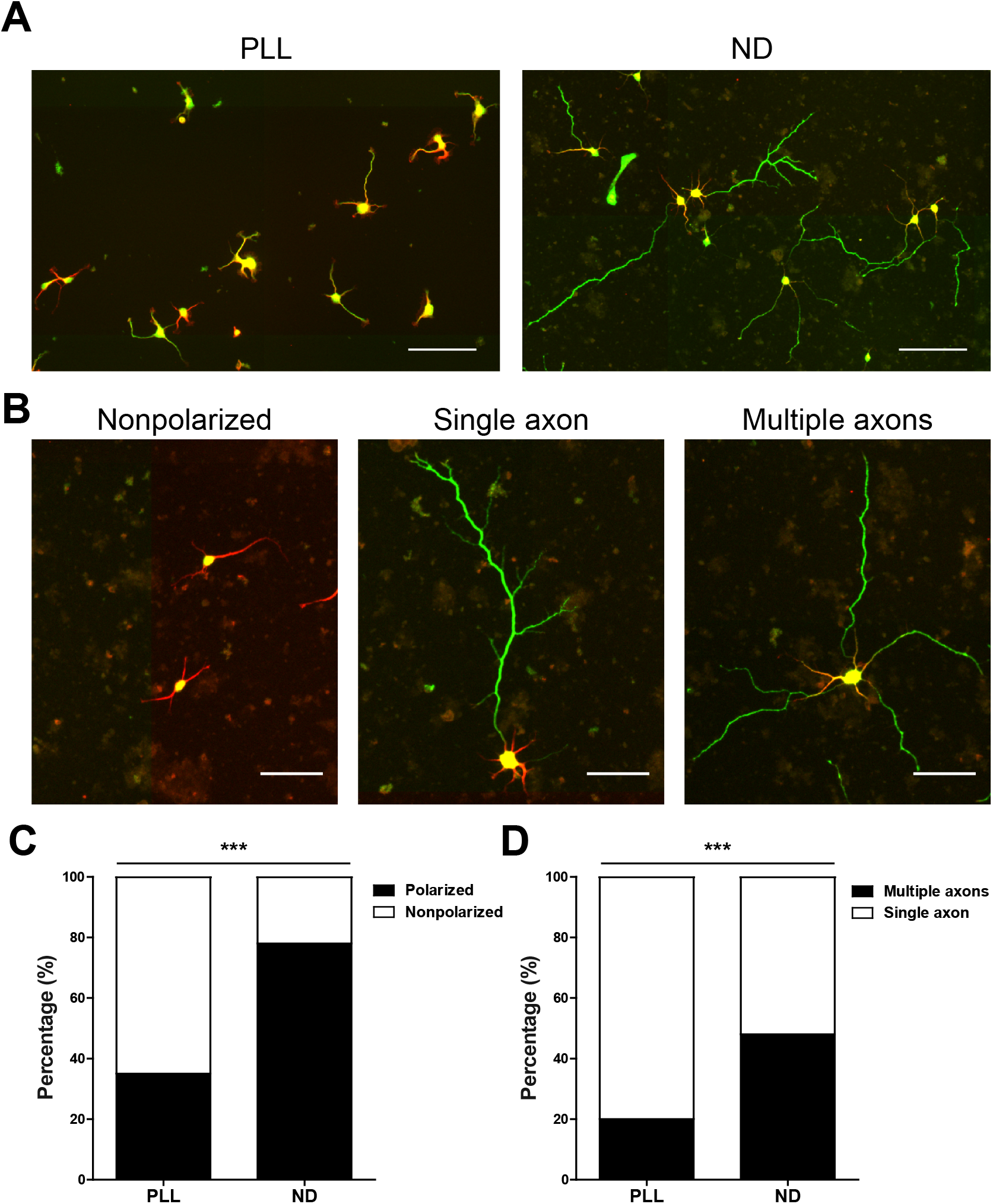
Nanodiamond surface accelerates neuronal polarization and induces the formation of multiple axons. (A) Representative images of dissociated hippocampal neurons cultured on PLL or ND surface for 40 hours. The density of ND coating was 37.5 μg/cm^2^. Cells were immunofluorescence stained with antibodies against MAP2 (red, dendrite-specific marker) and SMI312 (green, axon-specific marker). Scale bars represent 100 μm. (B) Representative images of nonpolarized, single axon, and multiple axons neurons on ND surface. Both dendrite (MAP2, red) and axon (SMI312, green) markers are shown. Scale bars represent 50 μm. (C) Percentage of polarized and nonpolarized neurons on PLL or ND surface. *** *p*<0.001, Fisher’s exact test. A total of 330 (PLL) and 279 (ND) neurons were quantified from 3 independent repeats. (D) Percentage of polarized neurons possessing single axon or multiple axons on PLL or ND surface. *** *p*<0.001, Fisher’s exact test. A total of 117 (PLL) and 218 (ND) polarized neurons were quantified from 3 independent repeats.

### Nanodiamond surface prevents soma aggregation

In addition to its effect on accelerating neuronal development, we also discovered that ND surface can reduce the aggregation of the neuronal cell bodies (somata). It has been shown that somata of mouse hippocampal neurons cultured on PLL surface in normoxic condition tend to cluster together (Chen et al., 2011; Peacock et al., 1979). In our hands, this clustering phenomenon is evident as early as 7DIV (**Figure 3A**). On the other hand, aggregation of somata is almost non-existent on the ND surface (**Figure 3A**). Automated neuronal morphology analysis on 7 DIV or 14 DIV hippocampal neurons shows that the average soma cluster area on ND surface is significantly smaller than that on PLL surface (**Figure 3B**, right), and the number of soma clusters on ND surface is significantly higher than that on PLL surface (**Figure 3B**, center). Both data demonstrate that somata are less aggregated on ND surface. In addition, the total area occupied by soma clusters of hippocampal neurons when cultured on ND surface does not significantly differ from that on PLL surface (**Figure 3B**, left), indicating that the attachment of neurons onto these two surfaces is similar. These results demonstrate that ND surface has an addition benefit of preventing somata aggregation during long-term culture.

**Figure 3.**
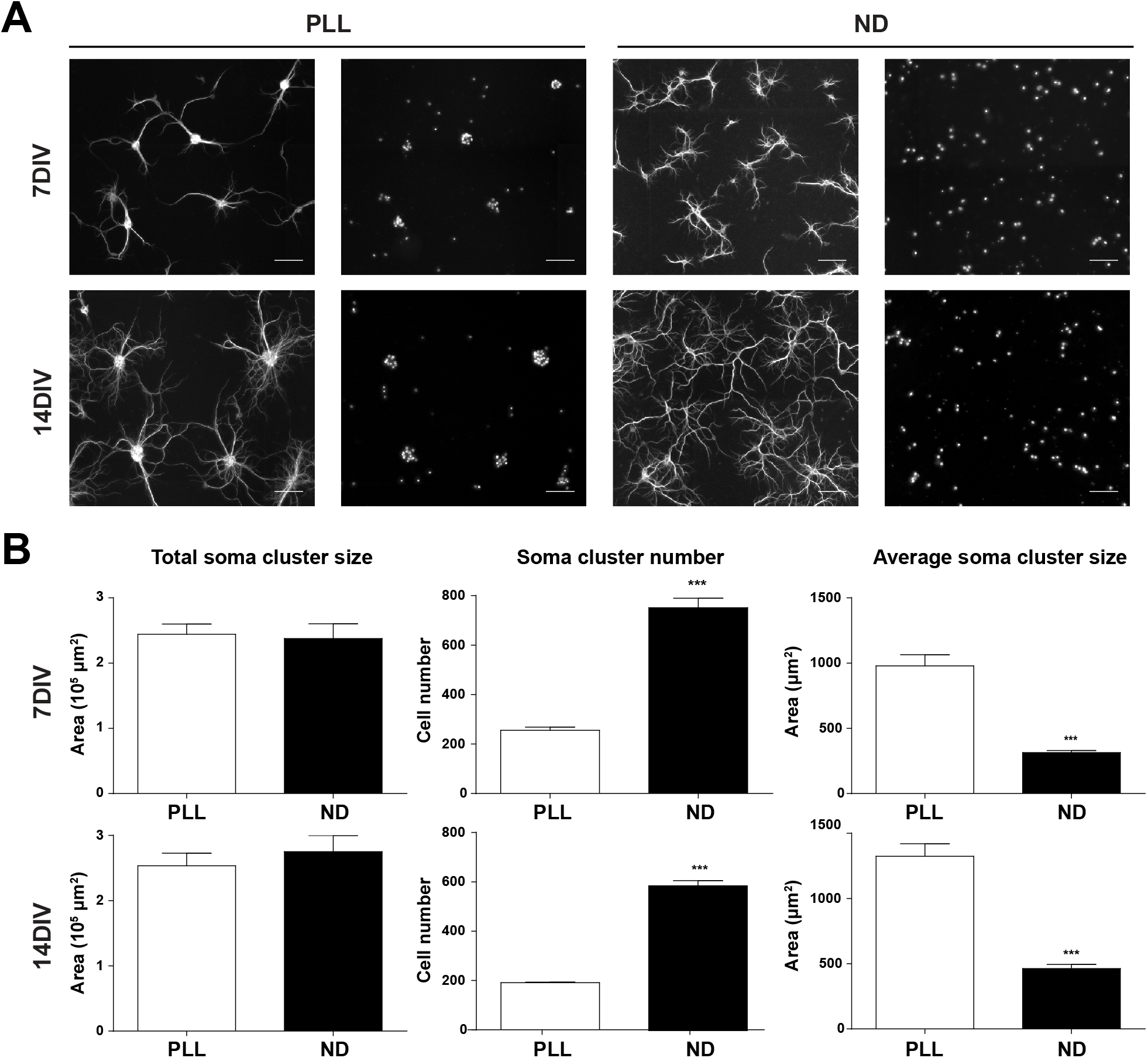
Nanodiamond surface prevents soma aggregation. (A) Representative images of 7DIV and 14DIV dissociated hippocampal neurons cultured on PLL or ND surface. The density of ND coating was 37.5 μg/cm^2^. Cells were immunofluorescence stained with the antibody against MAP2 (column 1 and 3) and a nuclear counterstain DAPI (column 2 and 4). Scale bars represent 100 μm. (B) Quantification of total soma cluster size (left), total soma cluster count (center), and average soma cluster size (right) of 7DIV (top) or 14DIV (bottom) hippocampal neurons cultured on PLL or ND surface. All bar graphs are expressed as mean and SEM from 3 independent repeats. *** *p*<0.001, two-tailed Student’s *t*-test.

### Nanodiamond surface reduces miR6236 expression

To better understand the biological response of neurons cultured on ND surface, RNAseq was utilized to examine the transcriptomic difference between neurons grown on PLL surface and those on ND surface. We reasoned that the transcriptomic changes would occur prior to the earliest morphological phenotype observed (i.e. neuronal polarization at 40 hours post-plating). As a result, neurons used for RNAseq were cultured on the PLL or ND surface for 36 hours and harvested for analysis. We first analyzed mRNAs known to enrich in the soma (GAPDH) (Briese et al., 2016) or along the neurite (β-actin) (Bassell et al., 1998) to confirm that ND surface does not prevent RNAs in certain cellular compartments from being collected (**Figure 4A-B**). Transcriptomes of hippocampal neurons collected from independent repeats are extremely consistent, demonstrating the high reproducibility of our sample preparations (**Figure 4C**). A comprehensive list of RNAs consistently detected (2 out of 3 independent experiments) from neurons cultured on two different surfaces is provided (**Supplementary Table 1**). The abundance of most highly expressed RNAs (<u>f</u>ragments <u>p</u>er <u>k</u>ilobase of transcript per <u>m</u>illion mapped reads, FPKM>500) are remarkably similar on PLL and ND surfaces (with a Pearson’s correlation coefficient of 0.995) except a single miRNA (mmu-mir-6236, hereafter referred to as miR6236) (**Figure 4D**). When different classes of RNAs are examined, miRNA is the only species that exhibits a significant abundance change (**Figure 4E**). miR6236 is by far the most abundant miRNA in dissociated hippocampal neurons on PLL or ND surface (**Figure 4F**). The overall level of miRNAs exhibits a 2.8-fold decrease in neurons cultured on ND surface, and miR6236 alone accounts for 97% of this reduction (**Figure 4F**). This transcriptomic analysis suggests that miR6236 is a candidate molecule conveying the surface topography to downstream post-transcriptional regulations.

**Figure 4.**
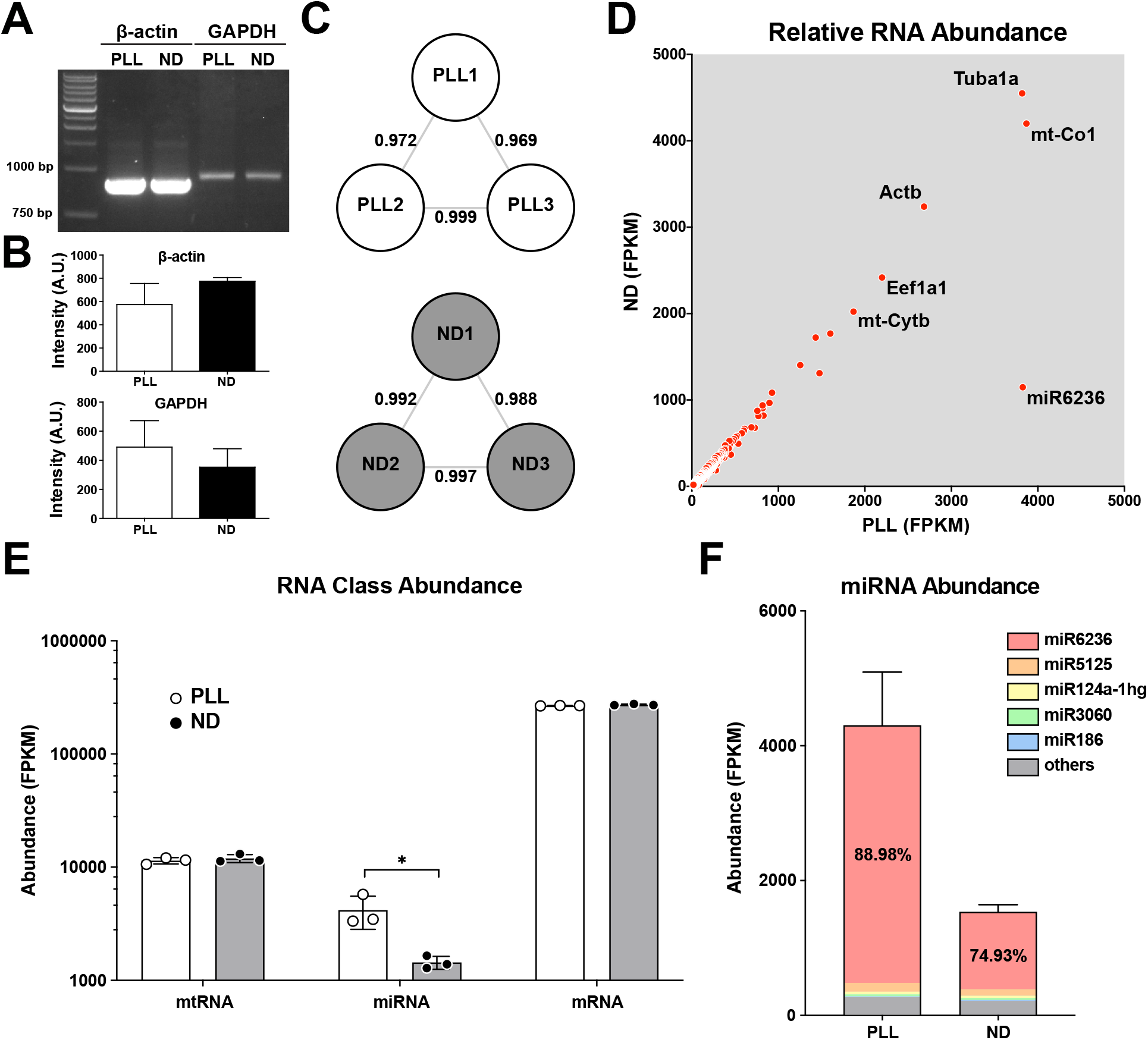
Nanodiamond surface reduces miR6236 expression. (A) RT-PCR examination of β-actin and GAPDH mRNA level in neurons cultured on PLL and ND surfaces. (B) Quantification of β-actin and GAPDH mRNA level. No statistically significant difference between PLL and ND samples were detected using two-tailed Student’s *t*-test. Both bar graphs are expressed as mean and SEM from 3 independent repeats. (C) Pearson’s correlation coefficient of all RNAs between independent repeats. The RNA quantity (fragments per kilobase of transcript per million mapped reads, FPKM) was used to calculate the Pearson’s correlation coefficient. (D) Relative RNA abundance between PLL and ND samples. The average quantity of the RNA from 3 independent repeats was used for graph creation. Predicted genes and pseudogenes were excluded from this graph. The top 5 most abundant RNAs in neurons cultured on PLL or ND surface are labeled. (E) The collective abundance of 3 classes of RNAs [mitochondrial RNA (mtRNA), microRNA (miRNA), mRNA] from neurons cultured on PLL or ND surface. Note the Y-axis is expressed in logarithmic scale. * *p*<0.05, two-tailed Student’s *t*-test. The bar graph is expressed as mean and SEM from 3 independent repeats. (F) miRNA abundance in neurons culture on PLL or ND surface. The top 5 most abundant miRNAs in neurons cultured on PLL or ND surface are labeled. miR6236 accounts for 88.98% and 74.93% of all miRNAs detected in PLL and ND sample, respectively.

### miR6236 depletion phenocopies the effect of nanodiamond surface

Since the RNAseq analysis revealed that miR6236 is the prime candidate in carrying out the biological response of the ND surface, we attempted to investigate this possibility. To deplete miRNA in primary neurons, miRNA sponge which specifically targets miR6236 was designed (Ebert et al., 2007). The miRNA sponge was constructed in the way that 12 copies of miR6236-binding sites are located at the 3’-UTR of the d2EGFP transcript expressed from the CMV promoter, while a mCherry transcript is expressed from a separate PGK promoter. This design enables the depletion of the target miRNA and allows the assessment of the target miRNA in the cell-of-interest by examining the ratio of d2EGFP/mCherry. A lower d2EGFP/mCherry ratio indicates a higher level of the target miRNA. As a control, a sponge based on the sequence of *CXCR4* (which is not complementary to any known mouse miRNA) was constructed. Dissociated neurons were transfected with the plasmid expressing either the control sponge or miR6236-targeting sponge just before plating. The lower d2EGFP/mCherry ratio in miR6236-targeting sponge expressing neurons indicated that miR6236 is present in neurons (**Figure 5A-B**). We then examined the effect of miR6236 depletion on neurite elongation and neuronal polarization by culturing neurons on PLL surface for 3 days and 40 hours, respectively. Interestingly, depleting miR6236 in hippocampal neurons on PLL surface phenocopies the effect of culturing neurons on ND surface. Firstly, neurons expressing miR6236 sponge extend longer neurites than those expressing the control sponge (**Figure 5A and C**). Secondly, a significantly larger proportion of neurons expressing miR6236 sponge become polarized than those expressing the control sponge (**Figure 5D-E**). Thirdly, miR6236 depletion results in a significantly higher proportion of neurons exhibiting multiple axons (**Figure 5F**). These results strongly suggest that miR6236 is the primary effector conveying the physical information provided by the ND surface.

**Figure 5.**
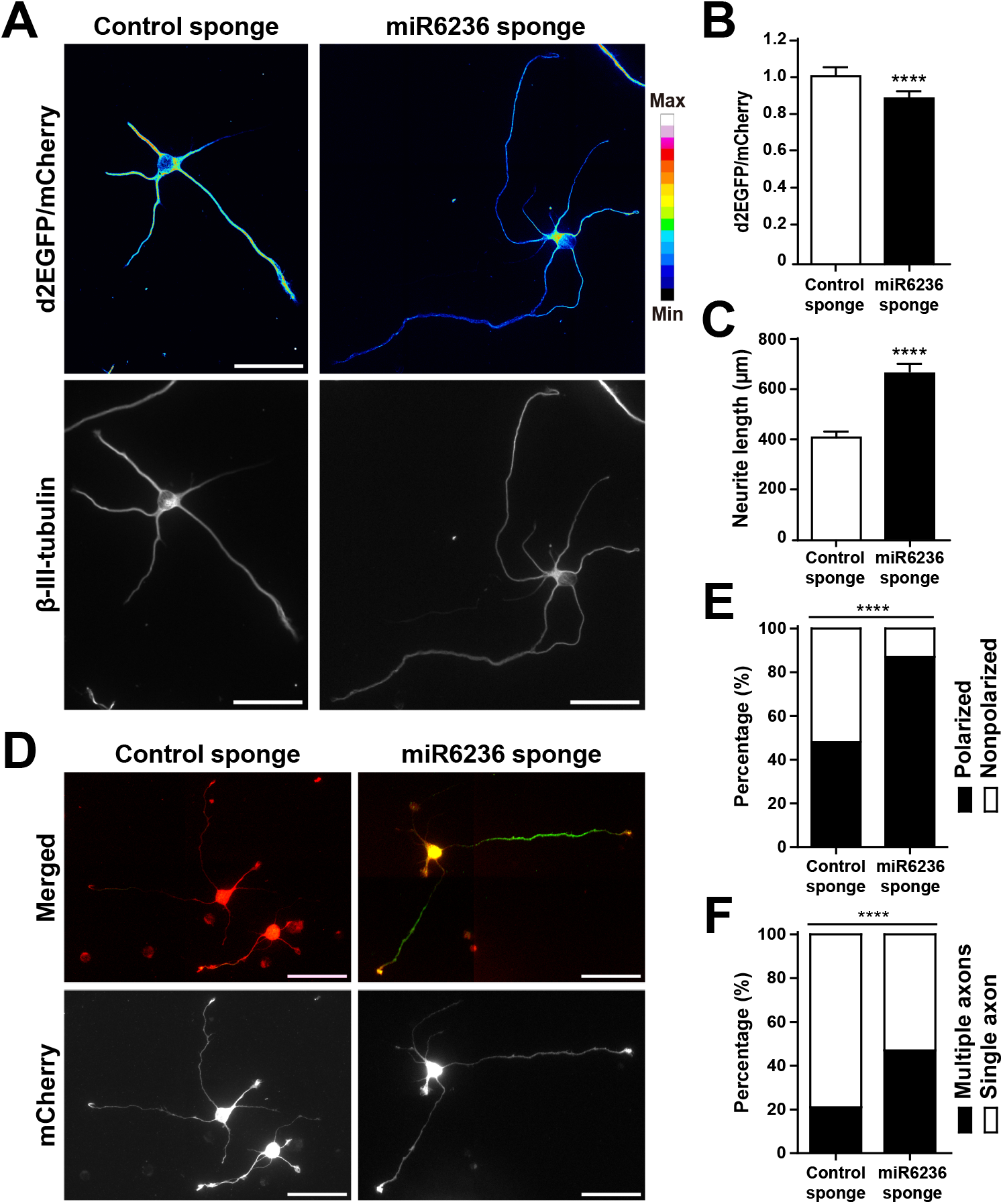
miR6236 depletion phenocopies the effect of nanodiamond coating. (A) Representative images of 3DIV dissociated hippocampal neurons expressing the control sponge (left) or miR6236-targeting sponge (right). Neurons are identified by immunofluorescence staining with the antibody against β-III-tubulin (bottom), and the ratio images of d2EGFP/mCherry are shown in pseudo-color (top). Scale bars present 50 μm. Quantification of d2EGFP/mCherry ratio (B) and total neurite length per neuron (C) in control or miR6236-targeting sponge expressing neurons at 3DIV. Both bar graphs are expressed as mean and SEM from 3 independent repeats. **** *p*<0.0001, two-tailed Student’s *t*-test. A total of 88 (control sponge) and 92 (miR6236 sponge) neurons were quantified from 3 independent repeats. (D) Representative images of dissociated hippocampal neurons transfected with the plasmid expressing the control sponge (left) or miR6236-targeting sponge (right) before plating and cultured for 40 hours. Both dendrite (MAP2, red) and axon (SMI312, green) markers are shown. Scale bars represent 50 μm. (E) Percentage of polarized and nonpolarized neurons. **** *p*<0.0001, Fisher’s exact test. A total of 218 (control sponge) and 217 (miR6236 sponge) neurons were quantified from 3 independent repeats. (F) Percentage of polarized neurons exhibiting single axon or multiple axons. **** *p*<0.0001, Fisher’s exact test. A total of 107 (control sponge) and 190 (miR6236 sponge) polarized neurons were quantified from 3 independent repeats.

In addition to the loss-of-function experiment mentioned above, a gain-of-function experiment was conducted to validate the effect of miR6236 on neuronal development. The plasmid expressing miR6236 or the control non-targeting miRNA was introduced into dissociated hippocampal neurons and cultured on ND surface. We first validated miR6236 expression plasmid using miR6236 sponge. As stated earlier, the d2EGFP/mCherry ratio of the sponge plasmid can be used to assess the level of the target miRNA. We therefore introduced the plasmid expressing miR6236 sponge and the plasmid expressing miR6236 into the neuron. As a control, the plasmid expressing miR6236 sponge and the plasmid expressing the control miRNA were introduced. Neurons expressing miR6236 sponge and miR6236 exhibit a significantly lower d2EGFP/mCherry ratio than those expressing miR6236 sponge and the control miRNA, confirming the function of the miR6236-expressing plasmid (**Figure S4**). We then tested whether miR6236 can inhibit neurite elongation and delay neuronal polarization. This is indeed the case as neurons expressing miR6236 possess shorter neurites (**Figure 6A-B**) and are less polarized (**Figure 6C-D)** than those expressing the control miRNA when cultured on ND surface. In addition, the proportion of multiple axon containing neurons is reduced when neurons are expressing miR6236 (**Figure 6E**). Taken together, these data indicate that ND surface suppresses the expression of miR6236, which negatively affects the morphological development of neurons.

**Figure 6.**
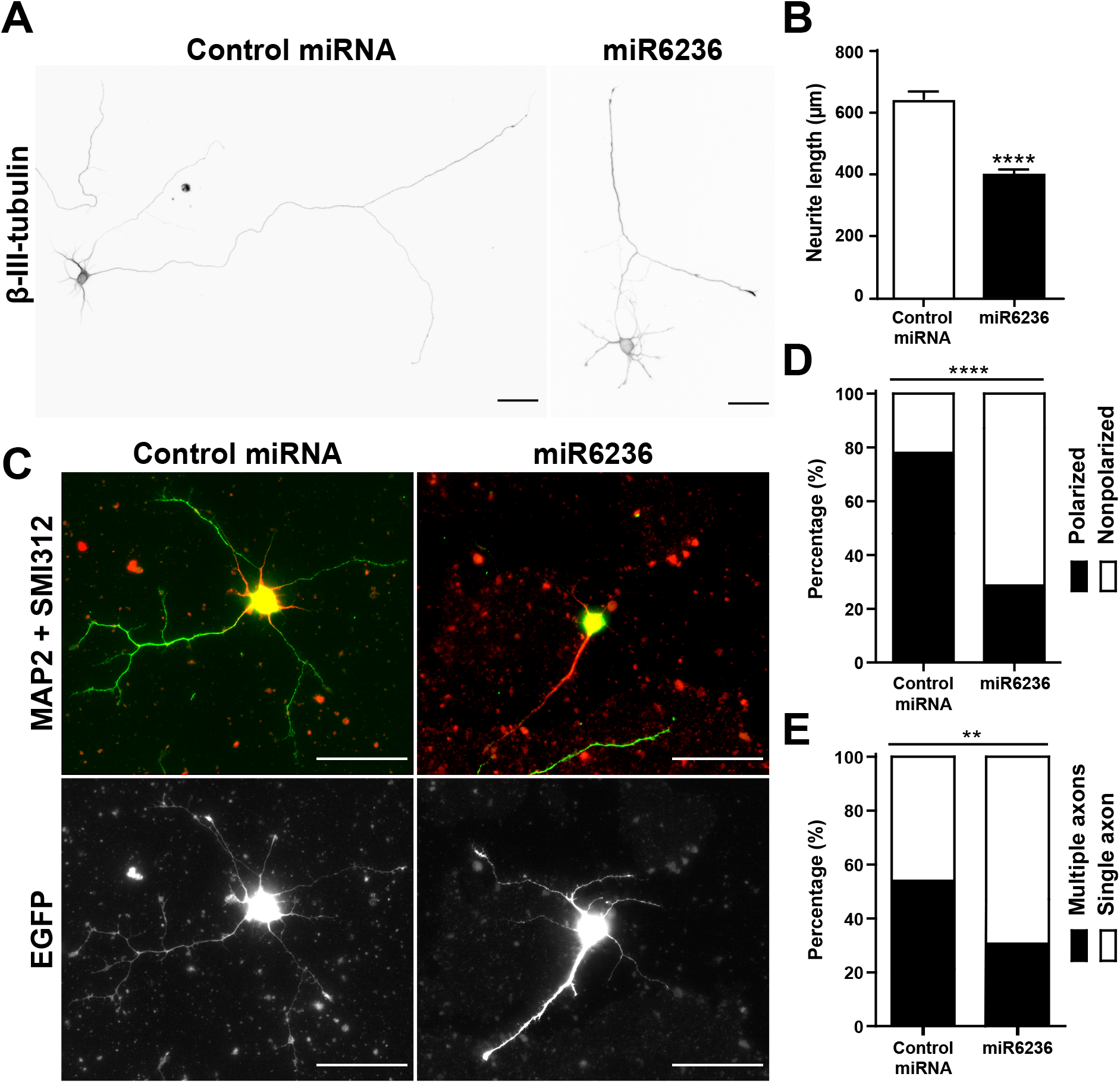
miR6236 overexpression reduces neurite elongation and neuronal polarization. (A) Representative images of dissociated hippocampal neurons transfected with the plasmid expressing the control miRNA (left) or miR6236 (right) before being cultured on ND surface for 3 days. Neurons are identified by immunofluorescence staining with the antibody against β-III-tubulin. Both scale bars present 50 μm. (B) Quantification of total neurite length per neuron. The bar graph is expressed as mean and SEM from 3 independent repeats. **** *p*<0.0001, two-tailed Student’s *t*-test. A total of 102 (control miRNA) and 104 (miR6236) neurons were quantified from 3 independent repeats. (C) Representative images of dissociated hippocampal neurons transfected with the plasmid expressing the control miRNA (left) or miR6236 (right) before being cultured on ND surface for 40 hours. Both dendrite (MAP2, red) and axon (SMI312, green) markers are shown. The extracellular clusters seen in the red and green channel came from the fluorescence of NDs. Scale bars represent 50 μm. (D) Percentage of polarized and nonpolarized neurons. **** *p*<0.0001, Fisher’s exact test. A total of 210 (control miRNA) and 217(miR6236) neurons were quantified from 3 independent repeats. (E) Percentage of polarized neurons exhibiting single axon or multiple axons in control miRNA or miR6236 group. ** *p*<0.01, Fisher’s exact test. A total of 158 (control miRNA) and 56 (miR6236) polarized neurons were quantified from 3 independent repeats.

### miR6236 depletion enhanced neuronal regeneration on inhibitory substrate

Our data thus far demonstrate that miR6236 negatively regulates neurite elongation in dissociated hippocampal neurons. Given that neurite elongation is tightly associated with neuroregeneration and *in vitro* neurite outgrowth screens have been used to successfully identify genes involved in neuroregeneration (Chandran et al., 2016; Moore et al., 2009), it is tempting to speculate that depleting miR6236 can promote post-injury neuroregeneration. To create the condition that mimics the post-injury environment *in vitro*, we plated dissociated hippocampal neurons on a surface coated with chondroitin sulfate proteoglycans (CSPGs). CSPGs are known to accumulate in the astroglial scar after CNS injury and inhibits the ability of neurons to regenerate (Yiu and He, 2006). We expect depleting miR6236 would increase the neurite length on CSPGs-coated surface. To deplete miR6236, the plasmid expressing miR6236 sponge was introduced into dissociated hippocampal neurons just before plating using electroporation. Transfected neurons were cultured on CSPGs-coated surface for 3 days and fixed to examine the effect of miR6236 depletion on neuroregeneration (**Figure 7A-B**). Consistent with our hypothesis, miR6236 sponge-expressing neurons on CSPGs-coated surface extend significantly longer neurites than those expressing the control sponge (**Figure 7C**). This result suggests that miR6236 depletion has the potential to promote regeneration in injured CNS neurons.

**Figure 7.**
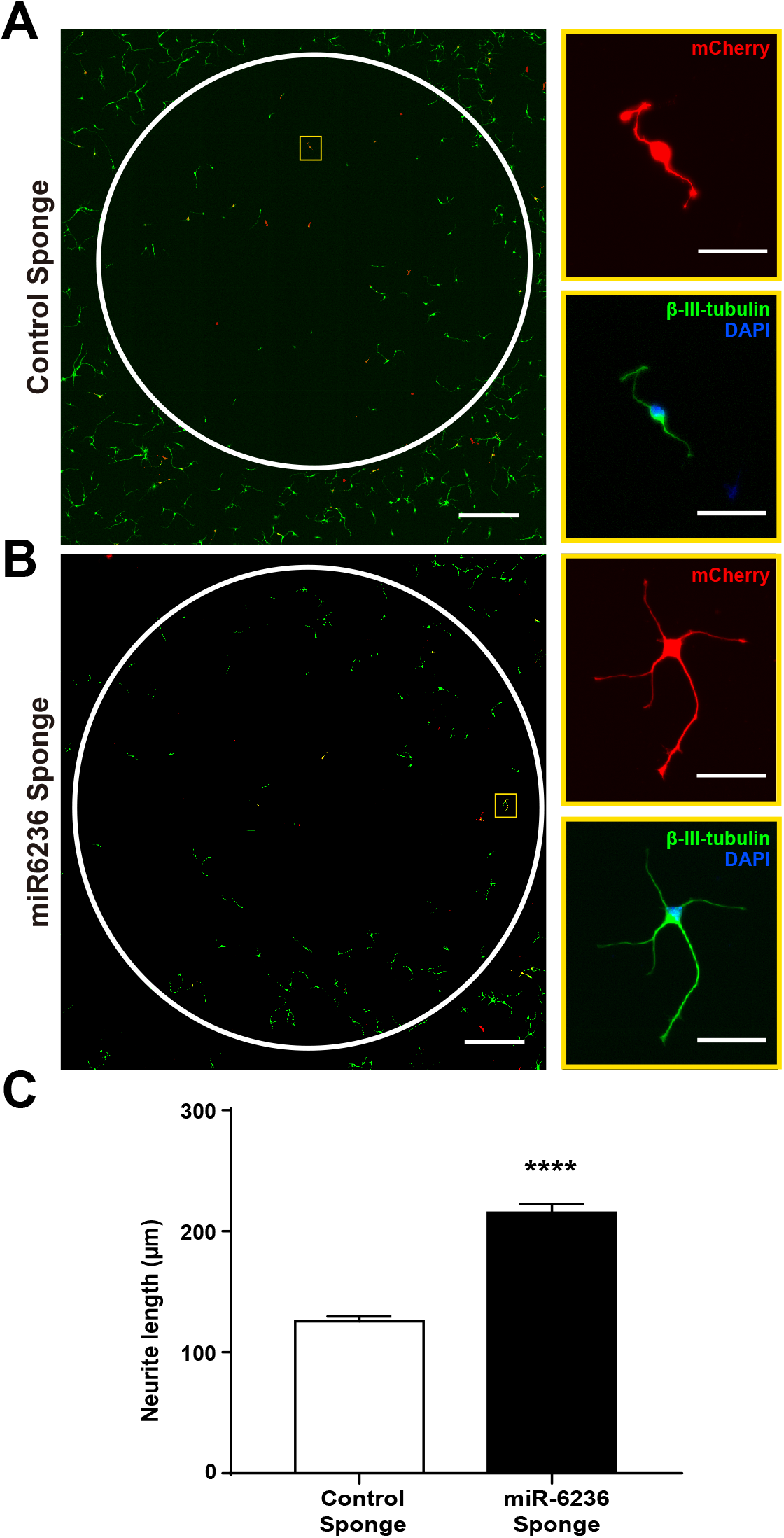
miR6236 depletion enhances neuronal regeneration on CSPGs-coated surface. (A-B) Representative images of 3DIV dissociated hippocampal neurons transfected with the plasmid expressing control sponge (A) or miR6236-targeting sponge (B), plated on CSPGs-coated surface (coated area marked by white circles), culture for 3 days, and fixed for staining. Fixed cells were immunofluorescence stained with antibody against β-III-tubulin (green). The magnified images on the right are from the yellow boxed areas in A-B, these images show the transfected neurons expressing mCherry (red) from the sponge-expressing plasmid, β-III-tubulin (green) and the nuclear counterstain DAPI (blue). Scale bars present 500 μm (left) and 50 μm (right). (C) Quantification of total neurite length per neuron inside CSPGs-coated area. **** *p*<0.0001, two-tailed Student’s *t*-test. The bar graph is expressed as mean and SEM. A total of 188 (control sponge) and 198 (miR6236 sponge) neurons were quantified from 3 independent repeats.

## Discussion

While biological mechanisms conveying chemical cues to a cellular response during neuronal development has been extensively studied (Ciani and Salinas, 2005; Guan and Rao, 2003; Huang and Reichardt, 2001; Tran et al., 2007), how physical cues are translated into cellular responses is largely unknown. In this study, we examined the nanotopography on neuronal development and how this physical cue is converted to the cellular response. The surface topography generated by NDs exhibits several distinct advantages for neurons. First, it promotes the attachment of primary neurons derived from both the central and the peripheral nervous systems. Second, ND surface enhances the morphogenetic processes of the attached neurons. These are consistent with the previous observation that ND surface promotes neuronal attachment and neurite outgrowth (Thalhammer et al., 2010). Third, ND surface prevents the aggregation of neuronal cell bodies during long-term culture. On the flip side, ND surface does possess adverse effect on cultured neurons. Hippocampal neurons cultured on ND surface developed multiple axon more frequently than those on PLL surface. While ND coating has been proposed as a viable option for therapeutic applications such as neurodegenerative interventions (Saraf et al., 2019), more in-depth analyses should be performed to understand its negative influence on the nervous system.

To identify the biological mechanism elicited by the ND surface, transcriptomes of neurons cultured on ND and PLL surface were compared. A single miRNA (miR6236) almost accounts for the entire transcriptomic change. miR6236 is by far the most abundant miRNA in young hippocampal neurons cultured on PLL surface. While still being the most abundant miRNA when neurons are grown on ND surface, its abundance exhibits a significant reduction. Depleting miR6236 in neurons cultured on PLL surface phenocopies the effects observed on ND surface, while overexpressing miR6236 in neurons cultured on ND surface suppresses the phenotypic effects. These observations suggest that effects caused by the ND surface are mainly mediated by miR6236. While miRNA expression have been linked to chemical cues during neuronal morphogenesis (Vo et al., 2005), little is known about the connection between miRNA and physical cues. As far as we know, this is the first report establishing a firm connection between the surface topography and a single miRNA during neuronal morphogenesis. It is possible that the developmental acceleration effect of hippocampal neurons caused by silica beads is also mediated by miR6236 (Kang et al., 2012b).

While the precise mechanism the ND surface leads to miRNA down-regulation is currently unknown, it is tempting to hypothesize what is at play. It has been shown that miRNAs can be specifically down-regulated at certain differentiation or developmental stages (Monticelli et al., 2005; Zhou et al., 2018). The abundance of miRNAs can be regulated during transcription or post-transcriptionally (Gebert and MacRae, 2019). miRNAs are transcribed by RNA polymerase II from introns or long non-coding RNAs (lncRNAs), and are subjected to transcription regulation (Ha and Kim, 2014). miR6236 localizes within a predicted gene which encodes a lncRNA (NR_168668). Since we did not perform lncRNA enrichment before our RNAseq analyses, the level of this lncRNA (NR_168668) cannot be determined. It will be interestingly to repeat the experiment using lncRNA enriched samples. Another mechanism to regulate miRNAs abundance is by degradation through target RNA-directed trimming, in which high complementarity between a miRNA and its target RNA leads to the 3’ to 5’ trimming of the miRNA (Ameres et al., 2010). It has recently been shown that miR-29 in the mouse cerebellum is specifically degraded via a near-perfect binding site located within the 3’ UTR of the *NREP* gene (Bitetti et al., 2018). One would assume that the miR6236 target RNA(s) that can lead to miR6236 degradation will exhibit elevated expression prior to miR6236 down-regulation.

In addition to discovering the regulation mechanism which leads to miR6236 down-regulation, it will also be interesting to identify the target RNAs of miR6236. Since the down-regulation of miR6236 leads to enhanced neuronal morphogenesis, it is reasonable to assume that proteins encoded by these target RNAs promote and/or accelerate neuronal morphogenesis. Using the RNAseq data collected at 36 hours post-plating, we compiled a list of RNAs that were up-regulated in neurons cultured on ND surface (**Supplementary Table 2**). It should be noted that RNAs whose expression is repressed by miR6236 may not exhibit significant increase at the time we collected samples for RNAseq. With this in mind, one of these up-regulated RNAs encodes myelin basic protein (MBP). MBP has been shown to cleave cell adhesion molecule L1 and promote neurite outgrowth in cerebellar neurons (Lutz et al., 2014). We are in the process of testing the level MBP in neurons cultured on ND or PLL surface and examining whether MBP can promote and/or accelerate neuronal morphogenesis in neurons.

When miR6236 was depleted from neurons cultured on the inhibitory substrate CSPGs, an enhanced neurite elongation can be observed. This indicates that miR6236 is involved in the inhibitory effect of CSPGs on post-injury neurite outgrowth. Given the emerging roles of miRNAs in spinal cord injury, traumatic brain injury, and stroke (Bhalala et al., 2013), it is a possibility that miR6236 is up-regulated following CNS injuries. It should be noted that most studies examining transcriptomic changes following the CNS injury were performed *in vivo* (Bhalala et al., 2012; De Biase et al., 2005; Hu et al., 2012; Liu et al., 2009; Redell et al., 2009; Redell et al., 2011; Strickland et al., 2011; Yunta et al., 2012). It is likely that the most prominent transcriptomic changes result from glial cells surrounding the injured neurons, and this may explain why miR6236 has never been discovered in those studies. It will be of great interest to perform microscopic image-based analysis on injured tissues to detect the change of miR6236 in neurons and glial cells.

## Materials and Methods

### Nanodiamond surface preparation

3~5 nm ND powder (1320JGY, NanoAmor, Katy, TX) was resuspended in sterile ddH_2_O to produce a 10 mg/mL stock solution. This stock solution was vortexed twice for 30 seconds, sonicated for 30 minutes, and transferred into a biosafety cabinet. The ND coating solution was made by diluting the 10 mg/mL stock solution with absolute ethanol to the indicated coating density. For example, 71.25 μL of the 10 mg/mL stock solution was mixed in 4 mL absolute ethanol, and 400 μL of this mixture was added into each well of a 24-well plate to produce the 37.5 μg/cm^2^ coating density. This ND-containing 24-well plate was then placed in an operating biosafety cabinet for 6 hours to allow the evaporation of ethanol.

### CSPGs surface preparation

Coverslips in 24-well plates were first coated with 100 μg/mL poly-L-lysine (PLL) overnight at room temperature. Next day, the coverslips were washed twice with sterile ddH_2_O. 6 drops of CSPGs (2.5 μg/mL diluted in 1× PBS, 2 μL per drop) were applied onto each coverslip in a 24-well plate and incubated overnight at 4°C. To help visualize CSPGs-coated area, Alexa Fluor 488-conjugated secondary antibodies (1:1000, Thermo Fisher Scientific, Waltham, MA) were added into the CSPGs solution.

### Neuron culture and transfection

All animal experimental procedures were approved by the Institutional Animal Care and Use Committee (IACUC) and in accordance with the Guide for the Care and Use of Laboratory Animals of National Chiao Tung University. Dissociated hippocampal neuron cultures were prepared as previously described with slight modification (Chen et al., 2017). Briefly, hippocampi from E17.5 mouse embryos were dissected, digested with trypsin-EDTA, and triturated. Dissociated neurons were seeded onto PLL-, ND-, or CSPGs-coated coverslips (500 cells/cm^2^ for examining neuronal polarization or 3×10^4^ cells/cm^2^ for other experiments) in the serum-containing neuronal plating medium (MEM supplemented with 2 mM L-glutamine, 5% fetal bovine serum, and 0.6% D-glucose). 4 hours later, the serum-containing neuronal plating medium was replaced by neuronal maintenance medium (neurobasal medium supplemented with 0.5 mM L-glutamine and 1× B27, for 3×10^4^ cells/cm^2^ density culture) or conditioned neuronal maintenance medium (for 500 cells/cm^2^ density culture). To obtain the conditioned neuronal maintenance medium, 2.3×10^6^ dissociated mouse cortical neurons were plated onto a 10 cm culture dish and incubated for 7 days. The cortical neuron culture medium was collected and used as the conditioned neuronal maintenance medium. 2×10^5^ dissociated neurons were transfected with Lonza Nucleofector using the Basic Neuron SCN Nucleofector kit (Lonza, Basel, Switzerland). Transfected neurons were incubated for indicating days and processed for subsequent analyses.

### Indirect immunofluorescence staining

Cells were fixed with 3.7% formaldehyde for 20 minutes at 37 °C and then washed three times with PBS. Fixed cells were permeabilized with 0.25% triton X-100 in PBS for 5 minutes at room temperature. After being washed with PBS, cells were blocked with 10% BSA in PBS for 30 minutes at 37 °C. Cells were then incubated for 1 hour at 37 °C with different primary antibodies: anti-β-III-tubulin (1:4000, 801202, Biolegend, San Diego, CA), anti-neurofilament SMI312 (1:1000, 837904, Biolegend, San Diego, CA), and anti-MAP2 (1:2000, AB5622, Millipore, Burlington, MA). After primary antibody incubation, cells were incubated with fluorophore-conjugated secondary antibodies (1:1000, Thermo Fisher Scientific, Waltham, MA). All antibodies were diluted in 2% BSA in PBS. Cells were washed three times with PBS before microscopy imaging.

### RNA purification, RT-PCR, and RNAseq

The hippocampal neuron RNA was extracted with Blood/Cell Total RNA Mini Kit (RB050, Geneaid, Taiwan) according to the manufacturer’s instructions. Briefly, 2×10^6^ dissociated mouse hippocampal neurons were cultured in a 10 cm culture dish coated with PLL or ND for 36 hours (RNAseq) or 48 hours (RT-PCR). Neuronal maintenance medium was then aspirated and washed with cold PBS twice. PBS was next replaced with 1 mL RB buffer mixed with 10 μL β-mercaptoethanol and incubated at room temperature for 5 minutes. Afterwards, cells were scraped off the dish using a cell scraper and subsequently transferred into a 1.5 mL microcentrifuge tube. 1 mL of 70% ethanol was added to the lysate and mixed vigorously. The mixture was then transferred into a RB column and centrifuged at 14,000 g for 1 minute. The flow through was discarded; and 400 μL Wash buffer was then added to the RB column and centrifuged at 14,000 g for 1 minute. 100 μL DNase I reaction buffer was added to the center of the RB column and incubated for 15 minutes. The RB column was next washed with 400 μL W1 buffer once and 600 μL Wash buffer twice. After the last washing step the column was centrifuged at 14,000 g for 3 minutes to dry up the residual ethanol. The dried RB column was placed onto a new 1.5 mL microcentrifuge tube and 50 μL of RNase-free water was added in the center of the column membrane. Finally, RB column was centrifuged at 14,000 g for 3 minutes to elute the total RNA.

Reverse transcription was performed using RevertAid first strand cDNA synthesis kit (K1621, Thermo Fisher Scientific, Waltham, MA) accordingly to the manufacturer’s instructions. PCR was performed using DreamTaq DNA polymerase (EP0702, Thermo Fisher Scientific, Waltham, MA) with primers targeting β-actin (5’-TGTGAACGGATTTGGCCGTATTGG-3’ and 5’-TGGAAGAGTGGGAGTTGCTGTTGA-3’) and GAPDH (5’-CCCCTGAACCCTAAGGCCA-3’ and 5’-CGGACTCATCGTACTCCTGC-3’).

For RNAseq analyses, all RNA sample preparation procedures were carried out according to the official Illumina protocol by Welgene Biotechnology Company (Welgene, Taiwan). SureSelect Strand-Specific RNA Library Preparation Kit (Agilent, Santa Clara, CA) was used for library construction followed by AMPure XP Beads size selection. The sequence was directly determined using Illumina’s sequencing-by-synthesis (SBS) technology. Sequencing data (FASTQ files) were generated by Welgene’s pipeline based on Illumina’s basecalling program bcl2fastq v2.2.0.

### Plasmids construction

For depleting miRNA, 12 copies of sponge sequences were inserted into the pLKO-AS7w.mCherry-CMV-d2EGFP plasmid (Academia Sinica RNAi core, Taiwan) in the 3’ UTR of the reporter gene d2GFP (destabilized green fluorescenct protein) under the CMV promoter. A separate mCherry gene under the PGK promoter was included as a transfection indicator. The sponge sequences used are: 5’ -CCTGACTAAAGCGACGGC- 3’ (miR6236-targeting) and 5’ -AAGTTTTCAGAAAGCTAACA- 3’ (*CXCR4*-targeting control).

For overexpressing miRNA, the following oligonucleotide sequences were used: 5’ -GCCGTCGCCGGCAGTCAGG- 3’ (miR6236) and 5’ -GCGCGCTATGTAGGATTCGTT- 3’ (non-targeting miRNA). The miR6236 and non-targeting miRNA sequences were inserted into the pSUPER-CAG-EGFP plasmid under the H1 promoter using BglII and HindIII (Chen et al., 2017).

### Image acquisition

For fluorescence microscopy, immunofluorescence stained images were acquired with a Nikon Eclipse-Ti inverted microscope equipped with a Photometrics CoolSNAP HQ2 camera, and Nikon NIS-Element imaging software. 10× 0.45 N.A, 20× 0.75 N.A, 60× 1.49 N.A Plan Apochromat objective lenses were used to collect fluorescence images. For scanning electron microscopy, ND-coated samples were mounted on a stub and coated with platinum and then observed in a field emission scanning electron microscope (JSM-6700F, JEOL, Tokyo, Japan).

### Image analysis

The ImageJ plugin NeurpholgyJ (Ho et al., 2011) was used for extracting and quantifying the total neurite length, the neurite width (an indicator for neurite bundling), and the total amount of somata in the β-III-tubulin stained images.

To quantify neuronal polarization, immunofluorescence staining with antibodies against the axon-specific neurofilaments (SMI312) and the dendrite-specific microtubule-associated protein (MAP2) was used. SMI312-positive and MAP2-negative neurites were recognized as axons. If a neuron possessed at least one axon, it was defined it as a polarized neuron. Polarized neurons were further categorized into two types: single axon neurons (which possess only a single axon) and multiple axons neurons (which possess 2 or more axons).

To measure the soma aggregation level of neurons, the ImageJ plugin NeurpholgyJ mentioned above was used for quantifying the soma/cluster count and soma/cluster area in the MAP2 (a neuron- and somatodendritic compartment-specific microtubule-associated protein) images. The soma/cluster count refers to the number of somata (or soma clusters) detected in the MAP2 image. If multiple somata were clustered together, the plugin could not separate them and counted them as one single object. The total soma/cluster area is the total area occupied by all somata (or soma clusters). The average soma/cluster area refers to the average area occupied by each soma (or soma cluster), and is calculated by dividing the total soma/cluster area by the soma/cluster count.

### Statistical analysis

All statistical analyses were performed in GraphPad Prism 7 with the indicated statistical methods.

## Acknowledgements

This work was supported by grants from Ministry of Science and Technology, Taiwan (MOST 105-2320-B-009-005-MY3 and MOST 108-2628-B-009-003) and “Center for Intelligent Drug Systems and Smart Bio-devices (IDS^2^B)” and “Smart Platform of Dynamic Systems Biology for Therapeutic Development” from The Featured Areas Research Center Program within the framework of the Higher Education Sprout Project by the Ministry of Education (MOE) in Taiwan.

