## Supplementary Materials for "miR6236, a microRNA suppressed by the anisotropic surface topography, regulates neuronal development and regeneration"

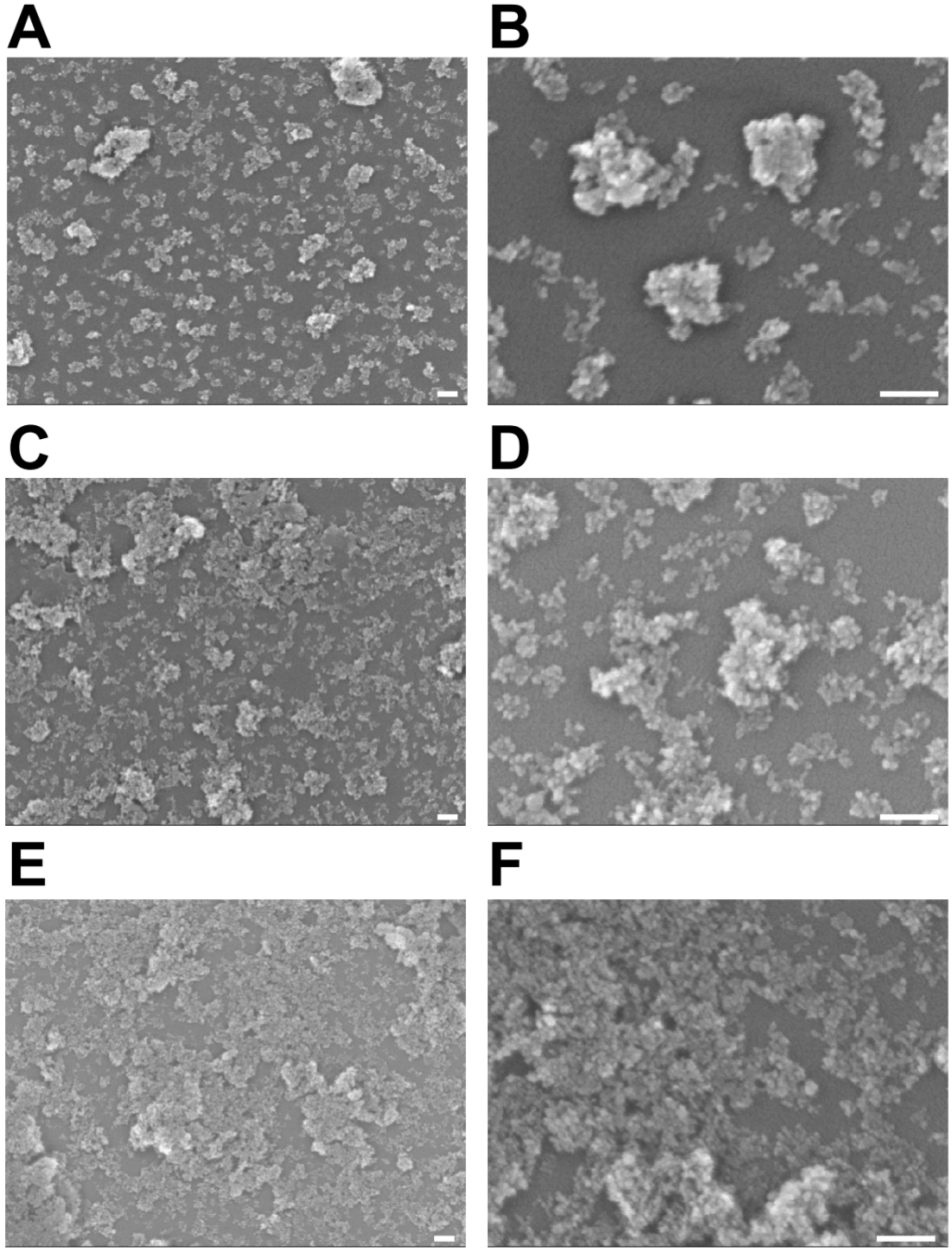


**Figure S1. Characterization of nanodiamond surfaces.**

Representative scanning electron microscopy images of the ND surface at various coating density. (A-B) ND coating at a density of 9.4 μg/cm$^2$. (C-D) ND coating at a density of 37.5 μg/cm$^2$. (E-F) ND coating at a density of 150 μg/cm$^2$. All scale bars represent 100 nm.

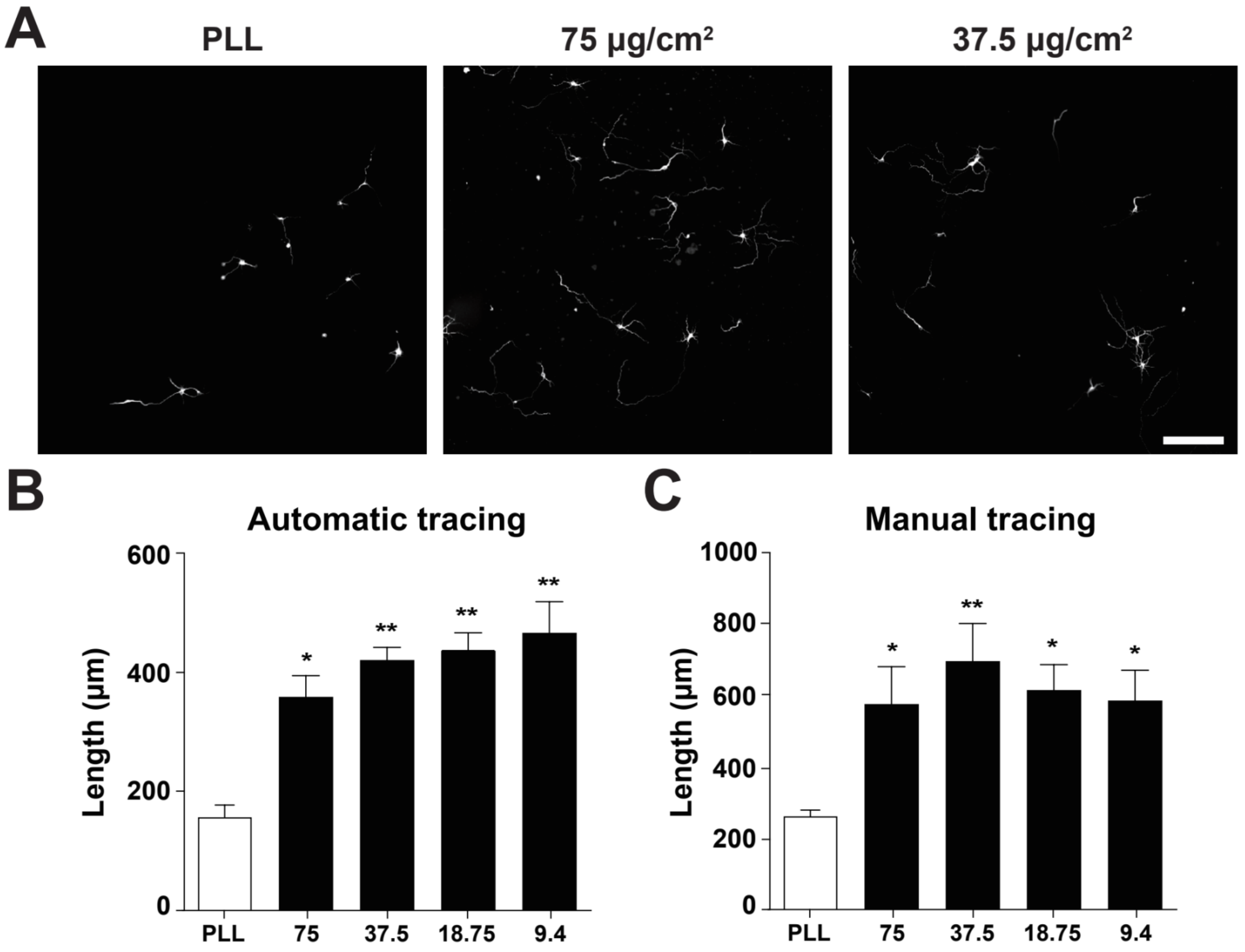


**Figure S2. Nanodiamond surface promotes neurite elongation.**

(A) Representative images of 3DIV dissociated hippocampal neurons seeded at $1\times10^3$ cells per well in a 96-well plate coated with PLL (left) or NDs (center and right). Neurons are identified by immunofluorescence staining with the antibody against β-III-tubulin. All images have the same scale and the scale bar represents 200 μm. Quantification of total length per neuron by the automatic tracing software NeurphologyJ (B) or by manual tracing (C). * $p<0.05$, ** $p<0.01$, one-way ANOVA followed by Dunnett's post-hoc analysis against the control group (PLL). Both bar graphs are expressed as mean and SEM from 3 independent repeats.

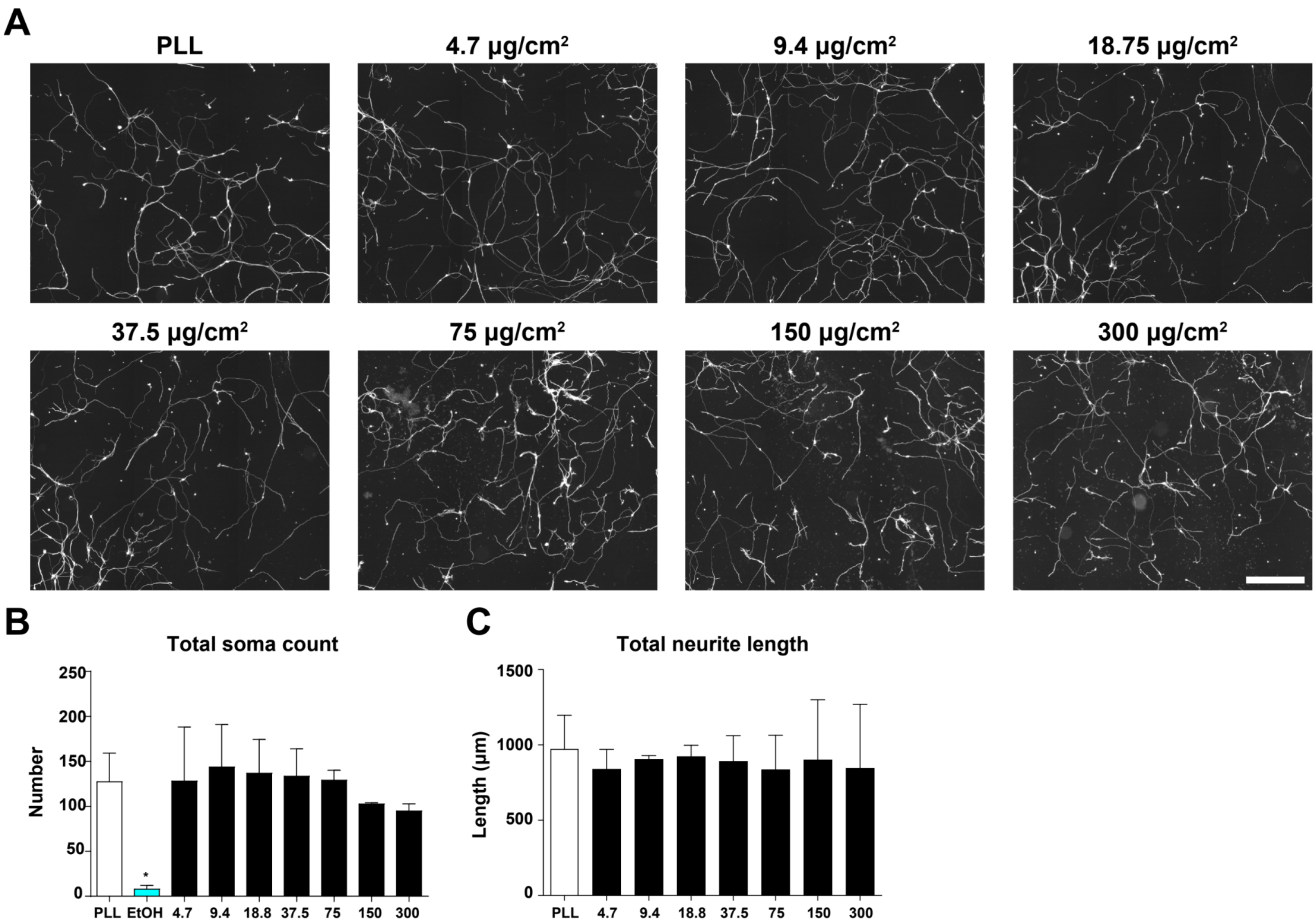


**Figure S3. Nanodiamond surface promotes the attachment of DRG neurons.**

(A) Representative images of 2DIV dissociated DRG neurons cultured on surfaces coated with various densities of NDs. Neurons are identified by immunofluorescence staining with the antibody against β-III-tubulin. All images have the same scale and the scale bar represents 500 µm. Quantifications of total soma count in a 13.91 $mm^2$ area (B) and total neurite length per neuron (C). * $p < 0.05$ from one-way ANOVA followed by Dunnett's post-hoc analysis against the control group (PLL). Both bar graphs are expressed as mean and SEM from 3 independent repeats.

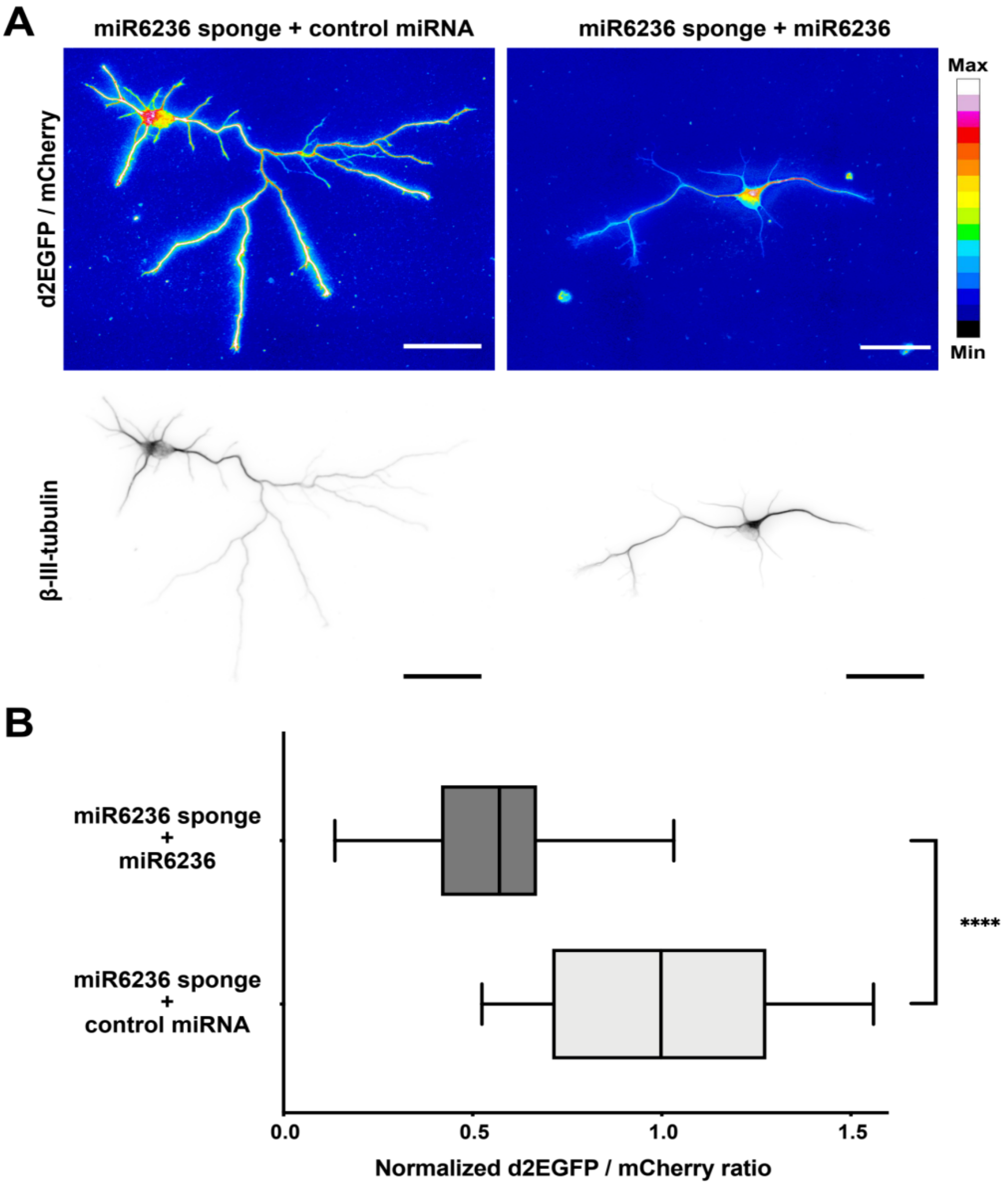


**Figure S4. Validating miR6236 overexpression plasmid in hippocampal neurons.**

To validate miR6236-expressing plasmids, both miR6236 sponge-expressing plasmid and miR6236-expressing plasmid were introduced into hippocampal neurons just before plating, incubated for 3 days, and fixed for d2EGFP/mCherry ratio quantification. As a control, plasmids expressing miR6236 sponge and control miRNA were introduced. (A) Representative pseudo-colored ratio images (top) or β-III-tubulin stained images (bottom) of 3DIV hippocampal neurons expressing the indicated plasmids. All scale bar present 50 µm. (B) Quantification of d2EGFP/mCherry ratio from 3 independent repeats. **** $p < 0.0001$, two-tailed Mann-Whitney test. The box plot shows first and third quartile with whiskers extending to 5-95 percentile.
